# Drinkable, liquid *in situ*-forming and tough hydrogels for gastrointestinal therapeutics

**DOI:** 10.1101/2022.12.15.520584

**Authors:** Gary W. Liu, Matthew J. Pickett, Johannes L. P. Kuosmanen, Keiko Ishida, Wiam A. M. Madani, Georgia N. White, Joshua Jenkins, Vivian R. Feig, Miguel Jimenez, Aaron Lopes, Joshua Morimoto, Nina Fitzgerald, Jaime H. Cheah, Christian K. Soule, Niora Fabian, Alison Hayward, Robert S. Langer, Giovanni Traverso

## Abstract

Tablets and capsules are a cornerstone of medicine, but these solid dosage forms can be challenging to swallow for geriatric and pediatric patients. While liquid formulations are easier to ingest, these formulations lack the capacity to localize therapeutics and excipients nor act as controlled release devices. To bridge the advantages of solid and liquid dosage forms, here we describe drug formulations based on liquid *in situ*-forming and tough (LIFT) hydrogels. Drug-loaded LIFT hydrogels are formed directly in the stomach through the sequential ingestion of a crosslinker solution of calcium and dithiol crosslinkers, followed by the ingestion of a drug-containing polymer solution of alginate and 4-arm poly(ethylene glycol)-maleimide. We show that LIFT hydrogels are mechanically tough and able to robustly form in the presence of complex gastric fluid and *in vivo* in rat and porcine stomachs. LIFT hydrogels are retained within the porcine stomach for up to 24 h, biocompatible, and safely cleared. These hydrogels deliver a total dose comparable to unencapsulated drug but with delayed and lower maximum drug plasma concentrations, providing a method for controlled release that may mitigate drug toxicity. Co-encapsulation of lactase as a model biologic drug and calcium carbonate mitigated gastric-mediated deactivation of encapsulated enzyme in rat and porcine models. We also demonstrate the potential of these hydrogels to encapsulate and protect a model therapeutic bacterium, *E. coli* Nissle 1917, against acid. LIFT hydrogels present a biocompatible means of tough, double-network hydrogel formation *in situ* in the gastric cavity, and may expand medication access for patients with difficulty swallowing.

## INTRODUCTION

The oral route provides a safe, rapid, and facile course for drug administration, and results in greater patient comfort and compliance compared to parenteral routes^1–5^. Due to advantages in stability, dose consistency, and the capacity to co-formulate with excipients, oral solid doses have become the predominant formulation: indeed, they consistently comprise ~50% of new FDA-approved drugs (fda.gov), and nearly 70% of Americans are on at least one prescription drug^6^. However, certain patient populations struggle with swallowing solids, complicating oral administration of solid drug forms. More than 50% of children are unable to swallow standard-sized pills or capsules^7^; among pediatric patients aged 0-6 years of age, 67.9% preferred liquid dosage forms compared to other forms^8^. Patients with dysphagia, or difficulty swallowing, similarly struggle with oral administration of solid drug forms^9^. In adults, prevalence of dysphagia can be as a high as 16%, and upwards of 37% have difficulty swallowing pills^9, 10^. This may cause patients to skip or modify (e.g., crush) their medications, which may result in altered pharmacokinetic profiles and even death^9, 11, 12^.

While liquid formulations are easier to ingest^2, 3^, they are susceptible to rapid dilution within the gastrointestinal tract and are unable to spatially localize drug with excipients^13^, which particularly challenge efforts to orally deliver biological drugs. A system capable of a programmed liquid-to-solid transition within the stomach could bridge the advantages of these two forms. Indeed, a solid matrix could facilitate spatial proximity of drug and excipients that modulate drug release or protect drug activity against the harsh gastric environment, and augment gastric residence of a drug depot. Efforts to develop liquid-to-solid systems have relied on drinkable hydrogel systems crosslinked by calcium. Xu *et. al* showed that orally administered calcium carbonate-loaded alginate solutions undergo gelation in the stomach due to acid-triggered release of Ca^2+^ ions and subsequent crosslinking of alginate^14^. Similarly, Kubo *et al*. reported *in situ* gastric gelation of gellan or alginate solutions mixed with complexed calcium^15^. Foster *et al.* demonstrated that oral administration of an alginate/karaya gum solution followed by a solution of CaCl_2_ resulted in gelation in the stomach^16^. However, these single-network hydrogels are mechanically weak and may not be able to withstand compressive forces within the stomach (up to 13 kPa)^17^, resulting in significant, irreversible deformation and potential breakage of the formulation within the gastrointestinal tract.

Here, we describe a new strategy to enable a drinkable, liquid *in situ*-forming and tough (LIFT) hydrogel, which comprises both ionic (calcium/alginate) and covalent (poly(ethylene glycol) (PEG)) polymer networks for enhanced toughness^18^. LIFT hydrogels undergo gelation after the liquid polymer solution containing alginate and functionalized PEG contacts the crosslinker solution within the stomach (**Fig. 1a, Fig. 1b**). We extensively characterize LIFT hydrogels after *ex vivo* formation in real gastric fluid and *in vivo* formation in rodent and large porcine models, and demonstrate that their capacity to form solids *in situ* enables these materials to act as a depot for controlled release of small molecules. Moreover, LIFT hydrogels can retain CaCO_3_ as an excipient and protect the activity of orally delivered enzymes and therapeutic bacteria from the low pH of the stomach in rodent and porcine models (**Fig. 1c**).

**Figure 1.**
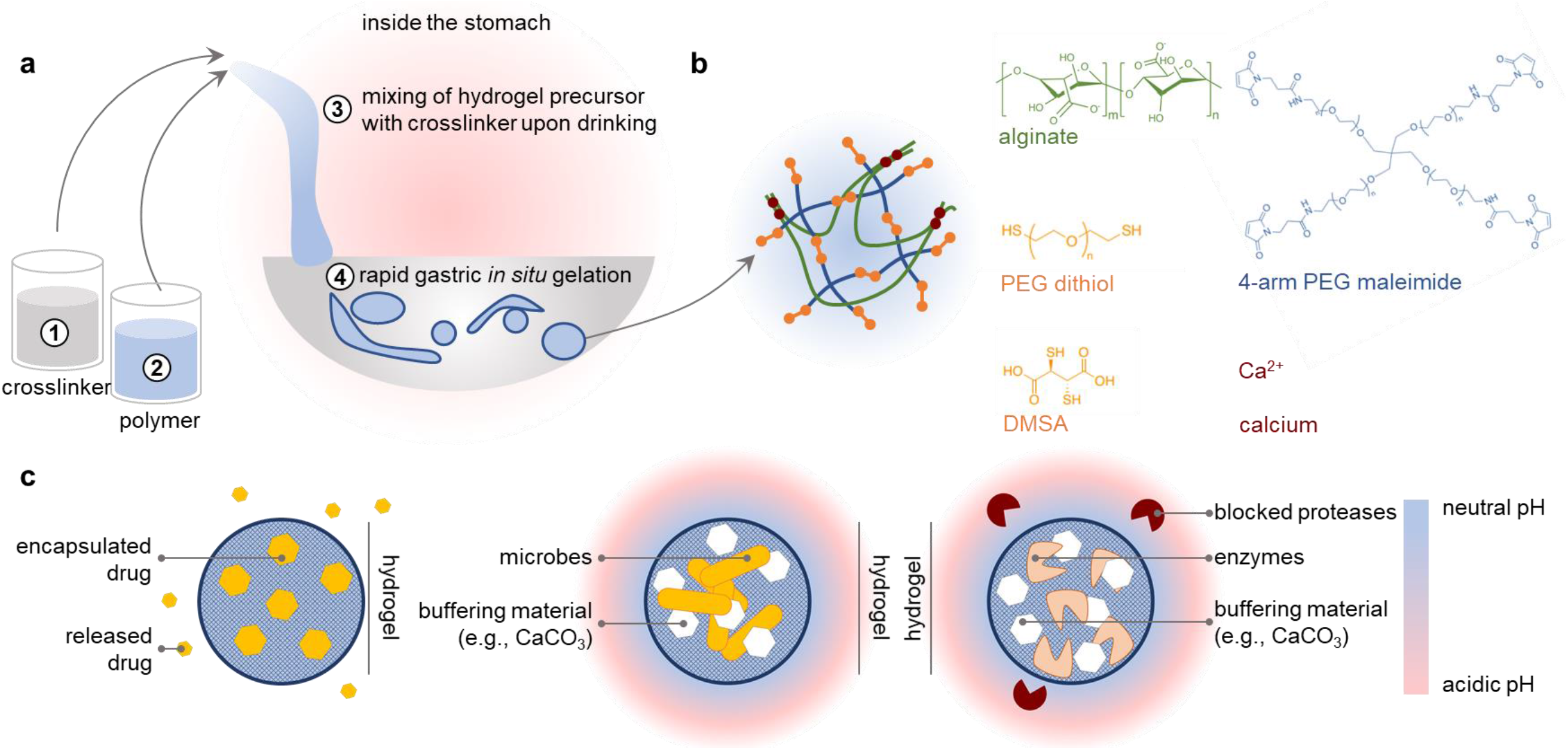
Overview of LIFT (liquid *in situ*-forming and tough) hydrogels. **A.** LIFT hydrogels form within the stomach after oral administration of (*1*) a crosslinker solution comprising CaCl_2_ and a dithiol-containing compound, followed by a (*2*) polymer solution comprising alginate and 4-arm PEG maleimide. These two solutions (*3*) mix within the stomach to form a tough double-network hydrogel (*4*) within the stomach. **B.**Schematic of the polymers and reagents used to facilitate crosslinking. Materials were selected due to their established safety profiles. Both a poly(ethylene glycol)-dithiol and dimercaptosuccinic acid (DMSA) were investigated as a dithiol crosslinker. **C.**Left: LIFT hydrogels may act as controlled release depot through encapsulation of water-insoluble drug that gradually dissolves and diffuses from the hydrogel. Middle, right: LIFT hydrogels enable co-encapsulation and co-localization of therapeutic microbes or enzymes and excipient (e.g., CaCO_3_) that modulate local pH and protect against proteases.

## RESULTS

Due to the relatively short residence times (<30 min) of liquids in the stomach and the complexity of gastric fluid^19, 20^, we prioritized the development of crosslinking chemistries that could rapidly and robustly crosslink two, interpenetrating polymer networks. Alginate is a well-studied, biocompatible polymer derived from algae with generally recognized as safe (GRAS) status; alginate polymers contain blocks of consecutive or alternating β-D-mannuronate and α-L-guluronate residues, the latter of which undergoes nearly instant crosslinking in the presence of calcium^21^. We utilized PEG for the second network due to the established safety profiles of ingested PEGs^22^, and initially considered three conventionally used crosslinking chemistries: NHS ester/amine, DBCO/azide, and maleimide/thiol. Due to the evolution of an uncharacterized and potentially toxic NHS leaving group during NHS ester/amine reaction and the slow kinetics (>1 h) of commercially available DBCO-/azide-functionalized PEGs (**Fig. S1**), we proceeded with development of a PEG network crosslinked by maleimide/thiol reaction. Advantages of this chemistry include its rapid reaction kinetics, mild reaction conditions, and biocompatibility^23^. To mitigate the risk of crosslinker toxicity, we focused our search on small molecular weight, FDA-approved or GRAS dithiol-containing molecules for rapid diffusion and crosslinking of maleimide-functionalized PEG. Dimercaptosuccinic acid (DMSA) was selected as a small molecule dithiol crosslinker due to its FDA approval status, extensive use history, and well-characterized safety profile in children and adults^24, 25^. A dithiol-terminated linear PEG (MW = 1000 Da) was also selected for evaluation. Therefore, our final concept comprises (*1*) ingestion of a crosslinker solution comprising calcium chloride and DMSA or PEG-dithiol, followed by (*2*)ingestion of a liquid polymer solution comprising alginate and 4-arm PEG-maleimide. Upon (*3*) mixing in the stomach, the liquid polymer solution undergoes crosslinking of both polymer networks and gelation to form (*4*) LIFT hydrogels (**Fig. 1a, Fig. 1b**).

We first asked whether LIFT hydrogels were capable of forming under short (20 min) time durations relevant to gastric residence of ingested liquids. To emulate *in vivo* formation conditions, a 0.5% w/v solution of alginate with 0, 5, and 10% w/v 4-arm PEG-maleimide was drop cast into a crosslinker solution (200 mM CaCl_2_/10 mM PEG-dithiol or DMSA) and then incubated for 10-20 min at 37 °C. The resulting hydrogels were mechanically characterized by compression testing. Notably, alginate hydrogels containing a crosslinked PEG network sustained significantly greater loads compared to alginate-only hydrogels (**Fig. 2a, Fig. S2a**). After 90% strain, LIFT hydrogels remained mostly spherical, whereas alginate-only hydrogels remained permanently deformed (flattened) (**Fig. 2b, Fig. S2b**). LIFT hydrogels were further mechanically characterized by cyclic compression testing. While LIFT hydrogels could sustain at least 5 cycles of 90% strain, alginate-only hydrogels remained permanently deformed after 1 cycle and were unable to sustain subsequent strains (**Fig. S3**). Due to the greater mechanical performance and easier manipulation of 0.5% alginate/5% w/v PEG-containing hydrogels compared to 10% w/v PEG-containing hydrogels, this composition was further characterized. To test the capacity of LIFT hydrogels to form *in vivo*, hydrogels were formed in fresh porcine gastric fluid at various dilutions in water. As a control, hydrogels were compared to LIFT or alginate-only hydrogels formed in the absence of gastric fluid. While gastric fluid attenuated the mechanical properties of LIFT hydrogels, these hydrogels were still mechanically tougher than alginate-only hydrogels formed under ideal conditions (**Fig. 2c**, **Fig. S2c**). LIFT hydrogel components were also tested for cytotoxicity in cultured human colon epithelial (Caco-2, HT-29), mouse liver (Hepa1-6), and monkey kidney (CV-1) cells. After 24 h of continuous incubation at relevant concentrations, no major causes of cytotoxicity were observed (**Fig. S4**). Collectively, these data demonstrate that LIFT hydrogels can form rapidly even in gastric fluid, the resulting hydrogels are mechanically tough, both DMSA and PEG-dithiol crosslinkers are capable of crosslinking the covalent PEG network, and that hydrogel components do not cause toxicity in cultured cells.

**Figure 2.**
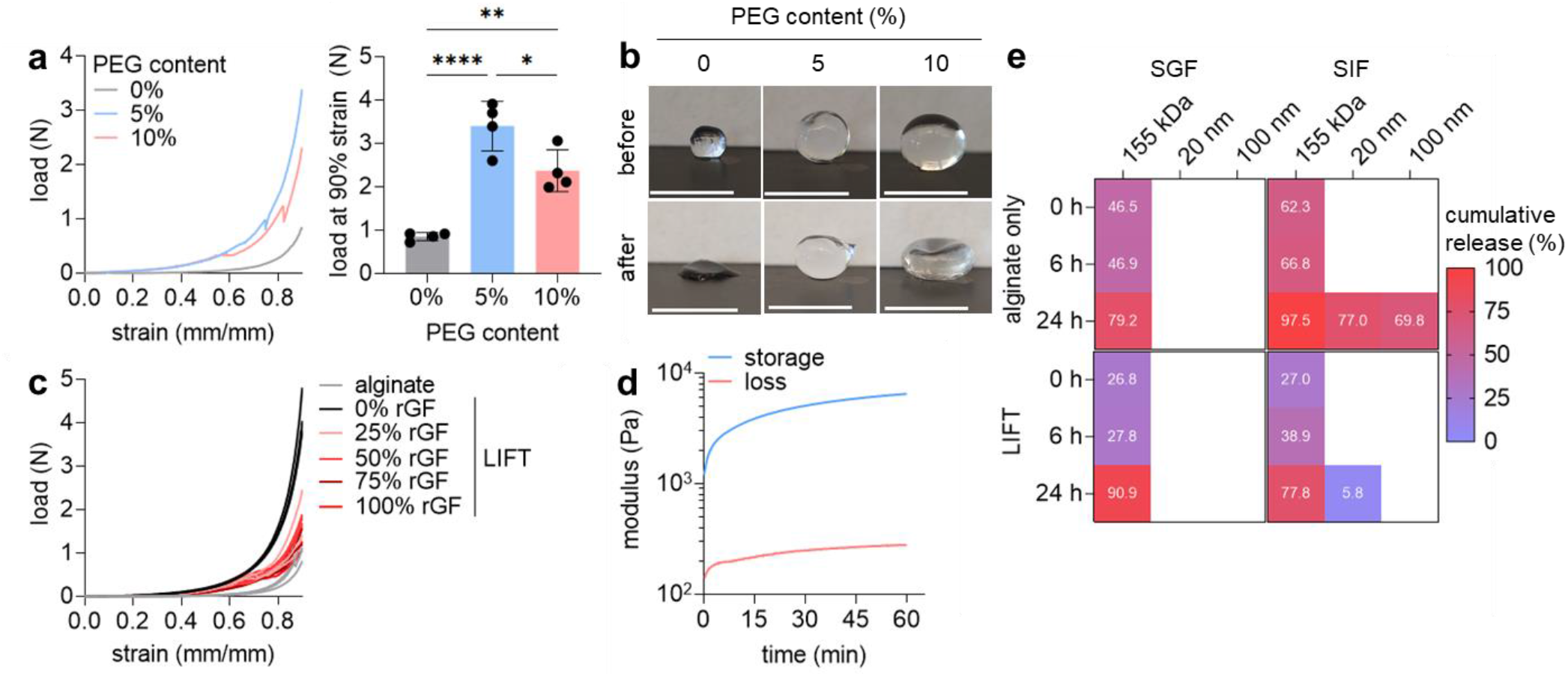
*In vitro* characterization of LIFT hydrogels. **A.** Left: load-strain curves of LIFT hydrogels comprising 0%, 5%, or 10% w/v 4-arm PEG-maleimide crosslinked in CaCl_2_/PEG-dithiol for 20 min, 37 °C, 50 RPM. Right: load at 90% of the different hydrogel compositions; *n* = 4 hydrogels were tested. **B.** Images of various compositions of hydrogels before and after 90% strain. Scale bar: 5 cm. **C.** Load-strain curves of LIFT hydrogels formed in various % v/v mixtures of gastric fluid in water containing CaCl_2_/PEG-dithiol. **D.** Gelation kinetics of LIFT hydrogels immersed in a crosslinker bath comprising CaCl_2_/PEG-dithiol at 37 °C, as characterized by rheology. **E.**Cumulative release of 155 kDa dextran and 20 or 100 nm nanoparticles from alginate and LIFT hydrogels. Hydrogels were incubated in SGF or SIF for the indicated time periods. \**p*-value < 0.05; \*\**p*-value < 0.01; \*\*\*\**p*-value < 0.0001. Bars represent mean ± standard deviation.

The kinetics of LIFT hydrogel formation were further studied by rheometry. To emulate rapid alginate crosslinking and to facilitate experimentation, hydrogels were first internally crosslinked with CaCO_3_ and glucono-δ-lactone and then analyzed within a bath of 200 mM CaCl_2_/10 mM PEG-dithiol solution. Notably, the sharpest increase in modulus occurred during the first 10-15 min (**Fig. 2d**), further supporting the feasibility of gastric crosslinking at timeframes relevant to liquid retention within the stomach (time of 50% emptying: 15-30 min)^19, 26^. LIFT hydrogels were then studied for their capacity to encapsulate therapeutic cargos of different length scales, using 155-kDa fluorescent dextran as a model macromolecule, and 20- or 100-nm fluorescent polystyrene nanoparticles as model control-release nanoparticles. The ability to co-encapsulate and retain cargos and excipients may facilitate protection of cargo function in the harsh gastrointestinal environment. LIFT or alginate-only hydrogels encapsulating these model cargoes were immersed in simulated gastric fluid (SGF, pH 1.77) or simulated intestinal fluid (SIF, pH 6.8), which were sampled at various timepoints. Neither hydrogels were able to detain dextran in either media (>75% release); however, LIFT hydrogels exhibited less nanoparticle release in SIF (<1-6%) compared to alginate-only hydrogels after 24 h (70-77%, **Fig. 2e**). This is consistent with prior reports describing the increased pore sizes and release of alginate hydrogels in alkaline environments^27, 28^. Therefore, LIFT hydrogels may be capable of retaining therapeutic cargoes at a variety of length scales due to greater stability at various pH ranges and/or smaller pore sizes.

LIFT hydrogels were then tested and characterized for formation, kinetics, and safety *in vivo*. Porcine models were tested due to the similarity in size of the gastrointestinal tract to that of humans^29^. First, the administration order of crosslinker (200 mM CaCl_2_/10 mM DMSA or PEG-dithiol) and hydrogel solution (0.5% alginate/5% w/v 4-arm PEG-maleimide) was varied. Pigs were anesthetized and administered solutions into the stomach *via* endoscope, and hydrogel structures were retrieved and studied 5-8 h afterwards. Hydrogels formed within the stomach cavity regardless of administration order. Administration of crosslinker solution first and then hydrogel solution resulted in the reproducible formation of noodle-like hydrogels within the stomach; conversely, administration of hydrogel solution first and then crosslinker resulted in larger bulk hydrogels (**Fig. 3a**, **Fig. S5, Fig. S6**). Accordingly, LIFT hydrogels were formed *in vivo* by first administration of the crosslinker solution followed by the hydrogel solution due to greater consistency in hydrogel architectures. LIFT hydrogels were then studied for their transit time *in vivo* through X-ray imaging of hydrogels containing 20% w/v barium sulfate. In general, LIFT hydrogels remained within the stomach up to 24 h after formation (**Fig. 3b**, **Fig. S6**); in comparison, liquids are emptied from the porcine stomach in 0.4-1.4 h across fasted and fed states^30^. No major changes in serum alkaline phosphatase, aspartate aminotransferase, alanine aminotransferase, blood urea nitrogen, or creatinine were observed up to 48 h after administration (**Table 1**). These data support that LIFT hydrogels and their components are safely cleared, do not cause obstructions, and do not cause toxicity. After formation in the gastric cavity, LIFT hydrogels were characterized for their mechanical properties by cyclic compression testing. Similar to *in vitro* experiments, LIFT hydrogels were tougher and able to sustain at least 5 cyclic 90% strains, whereas alginate-only hydrogels remained flattened after 1 cycle (**Fig. 3c-e**, **Fig. S7**). These findings highlight the capacity of the LIFT hydrogels to robustly form in the stomach after oral administration in a human-scale gastrointestinal tract.

**Figure 3:**
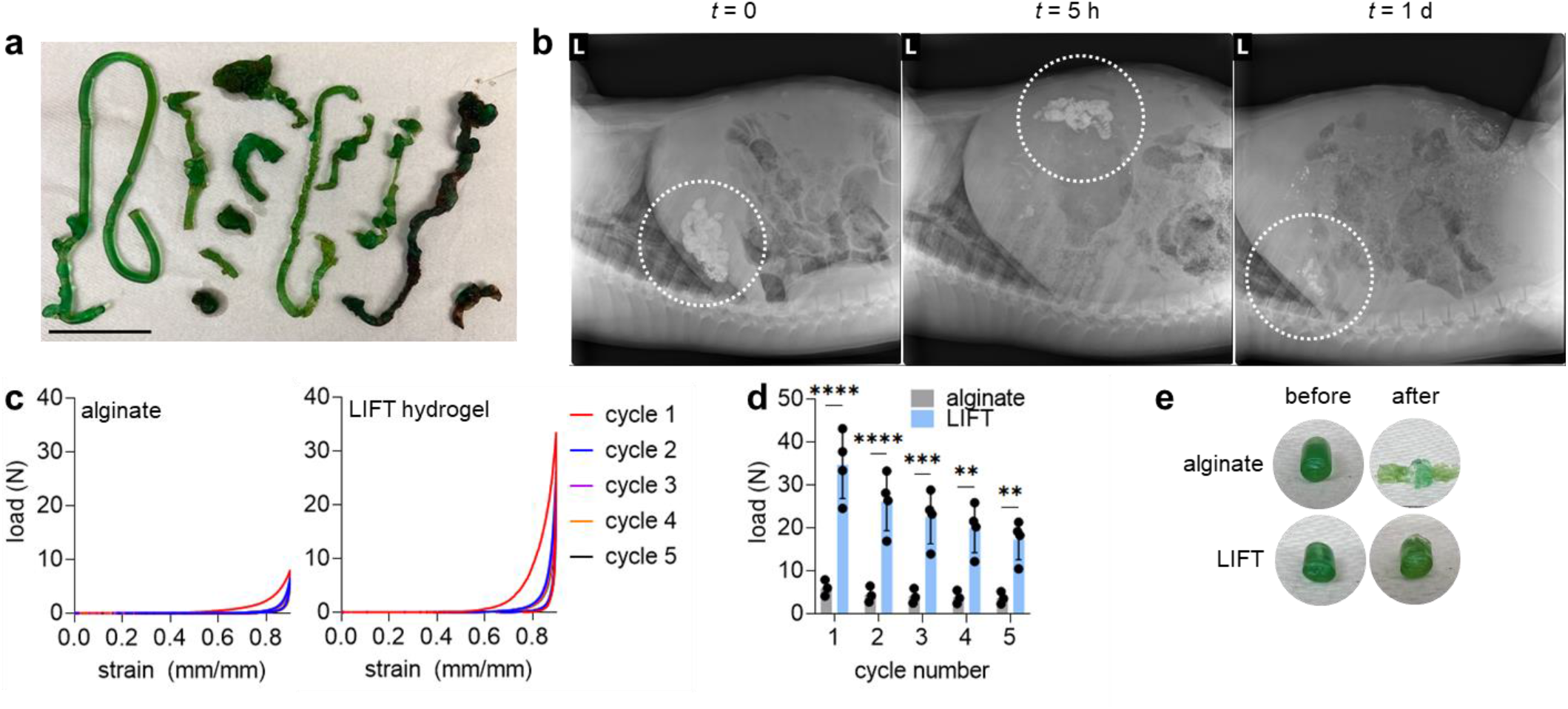
*In vivo* characterization of LIFT hydrogels. **A.** Hydrogel geometries after *in vivo* formation in pigs. LIFT hydrogels were formed by endoscopic administration of crosslinker solution (200 mM CaCl_2_/10 mM PEG-dithiol) followed by polymer solution (0.5% alginate/5% w/v 4-arm PEG-maleimide). Scale bar: 5 cm. **B.** X-ray imaging of LIFT hydrogels in pigs throughout time. Shown is representative of *n*= 3 independent experiments. **C.** Load-strain curves of alginate or LIFT hydrogels after retrieval from pig stomachs. Hydrogels were characterized by 5 cycles of 90% strain. **D.** Maximum loads experienced by alginate or LIFT hydrogels throughout 5 cycles of 90% strain. **E.** Images of retrieved alginate or LIFT hydrogels before and after 90% strain. \*\**p*-value < 0.01; \*\*\**p*-value < 0.001; \*\*\*\**p*-value < 0.0001. Bars represent mean ± standard deviation.

**Table 1.** Porcine blood chemistry after LIFT hydrogel administration.

| time | ALP<br>(U/L) | AST<br>(U/L) | ALT<br>(U/L) | BUN<br>(mg/dL) | creatinine<br>(mg/dL) |
| --- | --- | --- | --- | --- | --- |
| baseline | 131.0 ± 25.7 | 18.7 ± 4.0 | 41.7 ± 6.8 | 8.7 ± 0.6 | 1.1 ± 0.1 |
| 24 h | 132.3 ± 22.5 | 22.3 ± 17.2 | 41.7 ± 6.1 | 12.7 ± 4.5 | 1.2 ± 0.2 |
| 48 h | 115.0 ± 16.5 | 22.3 ± 5.5 | 38.7 ± 5.9 | 9.0 ± 1.7 | 1.1 ± 0.2 |

ALP, alkaline phosphatase; AST, aspartate aminotransferase; ALT, alanine transaminase; BUN, blood urea nitrogen. Baseline: sample was collected before LIFT hydrogel administration. An $n = 3$ pigs were tested.

Having established that LIFT hydrogels can form *in vivo*, we evaluated their capacity to encapsulate and modulate small molecule release and biologic drugs with excipients. We selected lumefantrine as a model small molecule drug because of its poor solubility in water and hence, would form a drug suspension that is encapsulated within the hydrogel after formation. Hydrogels were administered into the stomach of pigs using 200 mM CaCl_2_/10 mM DMSA as the crosslinker solution; lumefantrine was suspended in 0.5% alginate/5% w/v 4-arm PEG-maleimide LIFT polymer solution. Lumefantrine powder loaded in gelatin pills was used as a free drug control, and all pigs were dosed with 960 mg lumefantrine. Whereas free lumefantrine resulted in peak plasma concentrations at 5-7 h post-administration, hydrogel (alginate and LIFT) formulations resulted in peak plasma drug concentrations at ~24 h (**Fig. 4a**). The area under the curve (AUC) of released drug from free drug, alginate, LIFT hydrogel formulations was 14,873.5 ± 2,719.2, 7,568.4 ± 3,780.6, and 10,337.5 ± 3,849.7 ng·h/mL, respectively, and was not statistically different (**Fig. 4b**). While drug AUCs did not differ, the maximum observed drug concentration (C_max_) was significantly higher with free drug (901.2 ± 197.1 ng/mL) compared to alginate (283.8 ± 147.3 ng/mL) and LIFT (338.7 ± 112.6 ng/mL) hydrogel formulations (**Fig. 4c**). These data collectively support the capacity of LIFT hydrogels to deliver equivalent total doses of drug as free drug at lower plasma concentrations, which may reduce drug toxicity^9, 31^. We further envision that this system could be compatible with water-soluble drugs encapsulated within controlled-release particles suspended within the LIFT hydrogel, or mixed with drug powder to form tough *in situ* depots.

**Figure 4.**
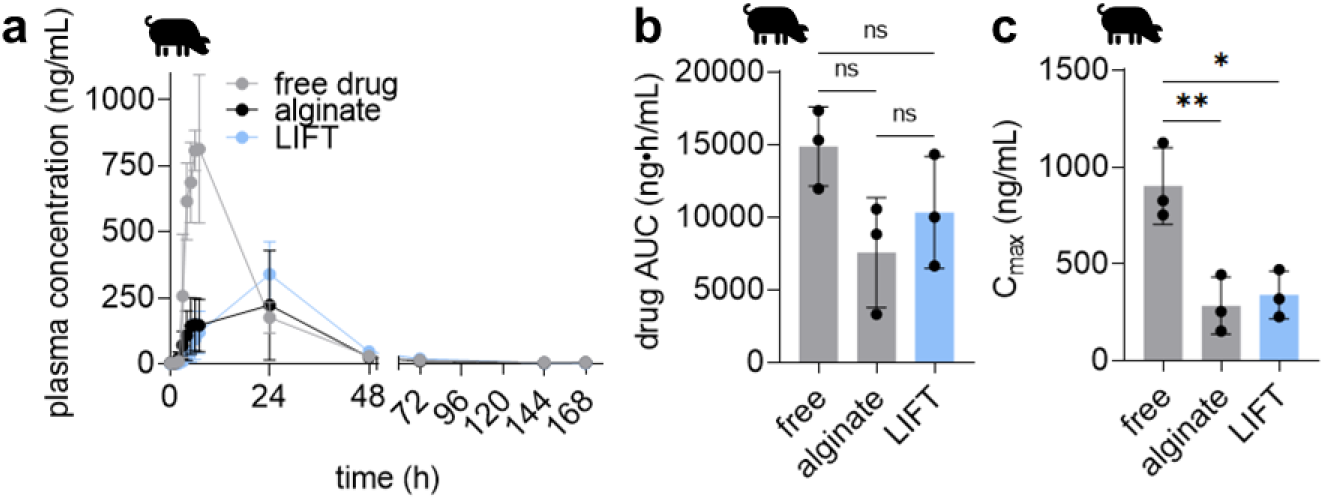
Pharmacokinetics of various oral lumefantrine formulations. **A.** Plasma lumefantrine concentration over time of free lumefantrine and lumefantrine (960 mg) encapsulated in alginate or LIFT hydrogel. For each treatment, *n* = 3 pigs were tested. **B.** Lumefantrine area under the curve (AUC) of each formulation. **C.** Maximum observed lumefantrine concentration (C_max_) of each formulation. \**p*-value < 0.05; \*\**p*-value < 0.01. Bars represent mean ± standard deviation.

We next evaluated the capacity of LIFT hydrogels for oral delivery of enzymes, which is challenging due to the acidic gastric fluid and proteases present within the gastrointestinal tract^32^. β-galactosidase (lactase) was selected as a model enzyme therapeutic due to the need for prolonged exogenous lactase activity in the stomach to mitigate the symptoms of lactose intolerance^33–36^. Indeed, lactase activity was found to be rapidly lost when incubated in SGF compared to PBS (**Fig. 5a**). Lactase was then encapsulated in alginate or LIFT hydrogels, along with calcium carbonate (CaCO_3_) as an excipient to neutralize the acidic gastric fluid. CaCO_3_ was selected because it is water-insoluble and therefore retainable within the hydrogels, and because of its GRAS status. Because the DMSA crosslinker attenuated lactase activity (**Fig. S8**), these LIFT hydrogels utilized the PEG-dithiol crosslinker. When challenged with SGF, CaCO_3_-containing hydrogels preserved lactase activity (**Fig. 5b**), underscoring the compatibility of LIFT hydrogels with enzymes. In addition to acidic gastric fluid, the gastrointestinal tract is rife with proteases that are capable of degrading enzymes. We next asked if the hydrogels were capable of protecting against trypsin as a model protease. Only LIFT hydrogels significantly preserved lactase activity compared to free lactase or lactase encapsulated in alginate hydrogels (**Fig. 5c**). Therefore, in addition to co-encapsulating CaCO_3_, LIFT hydrogels may exhibit additional barriers against exterior proteases due to the denser, dual polymer networks compared to alginate-only hydrogels^37–39^.

**Figure 5.**
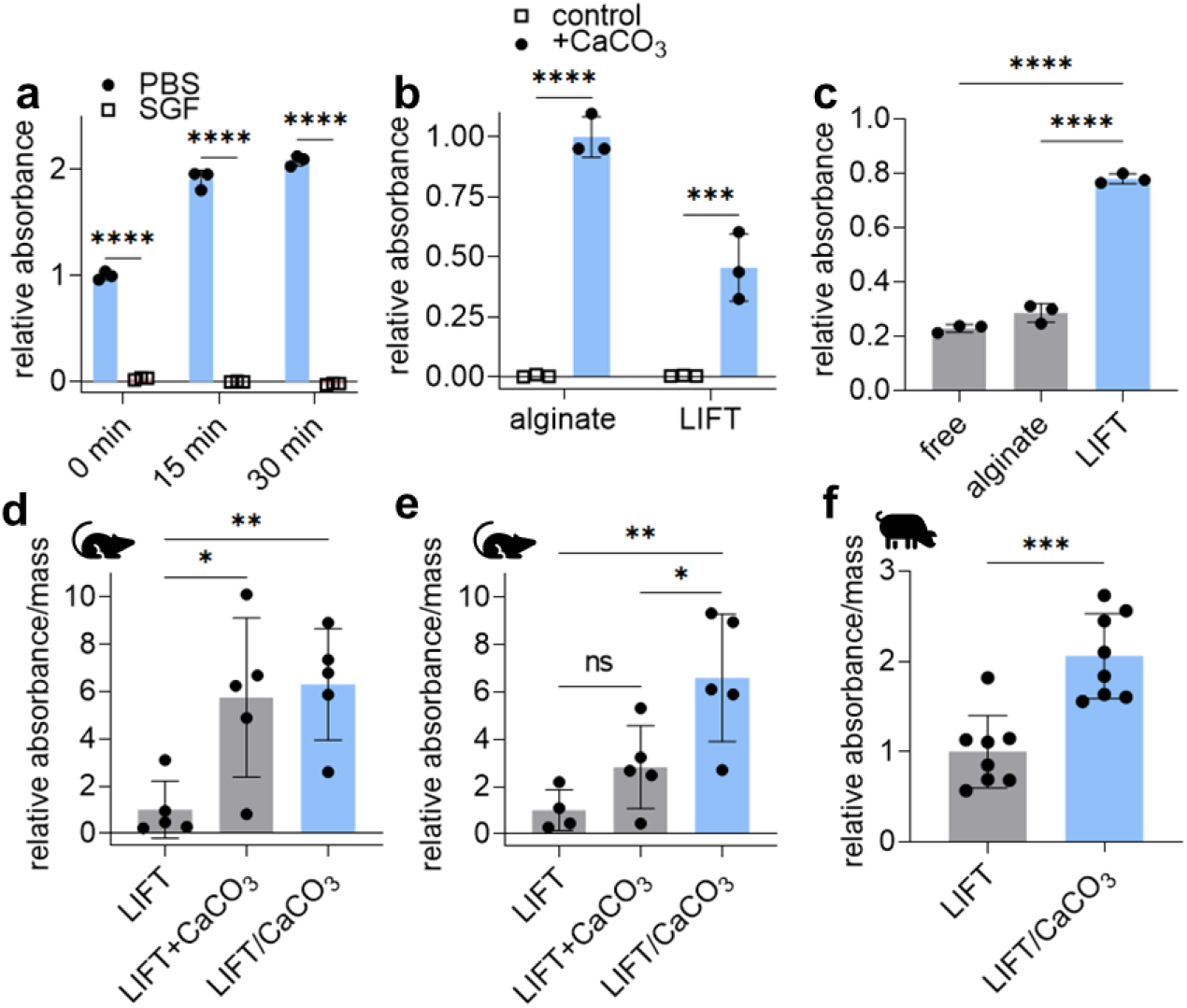
LIFT hydrogel co-encapsulation of CaCO_3_ protects lactase activity after oral delivery. **A.** Lactase activity, as measured by ONPG assay, after various incubations times in PBS or SGF at 37 °C. Absorbances were normalized to that of lactase incubated for 0 min in PBS. **B.** Lactase activity after hydrogel encapsulation with or without CaCO_3_ co-encapsulation and incubation in SGF for 1 h. Absorbances were normalized to that of alginate/CaCO_3_. **C.** Lactase activity of various treatment after trypsin treatment. Absorbances were normalized to that of treatment without trypsin. **D.** Activity of lactase encapsulated in LIFT hydrogels after 1 h in rat. CaCO_3_ was administered separately (LIFT+CaCO_3_) or co-encapsulated (LIFT/CaCO_3_). Absorbances were normalized by hydrogel mass; *n* = 5 rats were tested for each treatment. **E.** Activity of lactase encapsulated in LIFT hydrogels after 2 h in rat. Absorbances were normalized by hydrogel mass; *n* = 4 or 5 rats were tested for each treatment. **F.** Activity of lactase encapsulated in LIFT hydrogels after 6 h in pigs. Absorbances were normalized by hydrogel mass; shown is representative of *n* = 3 independent experiments. \**p*-value < 0.05; \*\**p*-value < 0.01; \*\*\**p*-value < 0.001; \*\*\*\**p*-value < 0.0001. Bars represent mean ± standard deviation.

LIFT hydrogels were then tested for their ability to protect lactase activity in rat and porcine models. We focused on analysis on LIFT instead of alginate hydrogels due to their capacity to protect encapsulated lactase against exogenous proteases. Similar to studies performed in pigs, rats were first administered the crosslinker solution by oral gavage immediately followed by the hydrogel solution containing lactase. Lactase was mixed in the hydrogel solution with or without CaCO_3_; as an additional control, CaCO_3_ was suspended in the crosslinker solution. Each animal was treated with a CaCO_3_ dose less than the maximum daily dose of 7-10 g/day (assuming a 75 kg human) established by manufacturers^40^. Therefore, these set of treatments test the effect of CaCO_3_ administered separately (LIFT+CaCO_3_) or co-encapsulated (LIFT/CaCO_3_). Oral gavage also resulted in robust hydrogel formation in rat stomachs (**Fig. S9**), and hydrogels were retrieved after *in vivo* incubation in stomachs and assayed for lactase activity. Notably, while separate and co-encapsulated CaCO_3_ significantly protected lactase activity after 1 h (**Fig. 5d**), only co-encapsulated CaCO_3_ protected lactase after 2 h (**Fig. 5e**). LIFT hydrogels with co-encapsulated CaCO_3_ resulted in 6-fold higher activity compared to control. The protective effect of the LIFT hydrogels co-encapsulating CaCO_3_ was also observed in porcine models after a 6 h *in vivo* incubation (2-fold higher than control, **Fig. 5f**), underscoring the advantage of oral systems capable of excipient co-encapsulation even in large animal models.

Another class of biologics of interest for oral delivery are therapeutic bacteria. Here, we utilized an engineered *E. coli* Nissle 1917 (EcN) that expresses both luciferase and luciferin; bioluminescence has been utilized to rapidly query bacterial viability and metabolism^41^. This particular strain was selected due to its safety and wide use as a chassis for synthetic biology therapeutics^42–44^. Oral delivery of bacteria is a recognized challenge and currently requires large solutions of sodium bicarbonate to buffer stomach pH^45^. Indeed, even short incubations of EcN in SGF pH 1.77 resulted in significant decreases in bioluminescence (**Fig. 6a**). Similar to lactase, EcN also exhibited sensitivity to DMSA crosslinker (data not shown); therefore, PEG-dithiol was used as the dithiol crosslinker in subsequent studies. We tested the capacity of LIFT hydrogels to protect bacterial activity in porcine models after 6-8 h incubation in the stomach. While bacteria encapsulated in LIFT/CaCO_3_ hydrogels exhibited greater average bioluminescence than those in LIFT hydrogels, this difference did not reach statistical significance (**Fig. S10**). Notably, bacteria were metabolically active after retrieval and able to proliferate after hydrogels were incubated in growth media (data not shown), suggesting that LIFT hydrogels are capable of sustaining bacterial viability *in vivo*.

**Figure 6.**
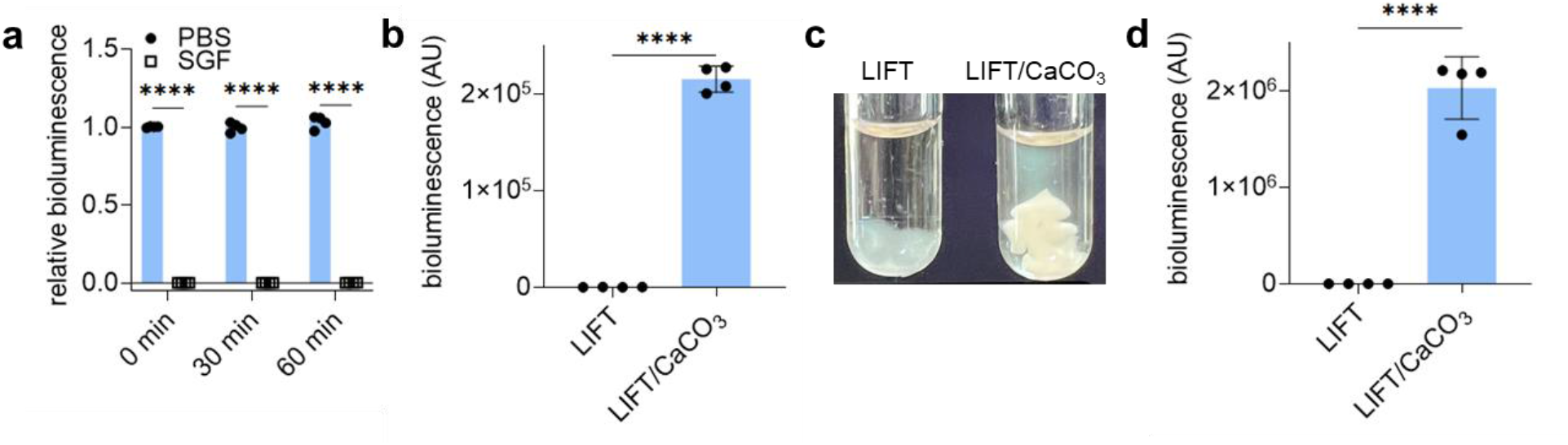
LIFT hydrogel co-encapsulation of CaCO_3_ protects bacterial activity. **A.** Bioluminescence of luciferase-expressing *E. coli* Nissle 1917 bacteria after various incubations times in PBS or SGF at 37 °C. Bioluminescence was normalized to that of bacteria incubated in PBS at each time point. **B.** Bioluminescence of bacteria encapsulated in LIFT hydrogels with or without CaCO_3_ and incubated in SGF for 3 h. **C.** Representative images of culture tubes containing LB culture medium after bacteria-containing LIFT hydrogels were challenged with SGF and then incubated in the culture tubes for 4 h at 37 °C. Cloudiness of medium indicates viable bacteria remain inside the hydrogel. **D.** Bioluminescence quantification of media supernatants from panel C. \*\*\*\**p*-value < 0.0001. Bars represent mean ± standard deviation.

The relatively high pH values of pig gastric fluid (pH > 5) may not be sufficient to reduce bacterial viability^46^. We reasoned that in terms of acidity, *in vitro* studies may provide a more aggressive challenge than *in vivo* treatment in pigs that is physiologically relevant to human gastric fluid (pH 1.4-2.1 in the fasted state)^47^. Therefore, we asked if LIFT hydrogels were capable of protecting bacterial bioluminescence upon encapsulation with or without CaCO_3_ and after challenge with SGF pH 1.77 for 3 h. Only co-encapsulation with CaCO_3_ resulted in bacterial bioluminescence compared to LIFT hydrogel control (**Fig. 6b**). To further confirm that bacteria were indeed viable and capable of growth, hydrogels were incubated in growth media after SGF challenge. While the media of LIFT hydrogels remained clear, the media of LIFT/CaCO_3_ hydrogels appeared turbid, suggesting bacterial proliferation (**Fig. 6c**). The supernatant was quantified for bioluminescence to confirm bacterial viability and metabolism; only supernatant from LIFT/CaCO_3_ hydrogels exhibited bioluminescence (**Fig. 6d**). Therefore, LIFT hydrogels are capable of supporting bacterial viability and protection against acid challenge when loaded with CaCO_3_. Given that acid secretion can vary 40-71 mmol/h (interquartile range) in humans and the recommended maximum recommended dose of CaCO_3_ (10 g/day)^40, 48^, these systems could potentially support the viability of bacteria in the stomach for at least 3-5 h.

## DISCUSSION

Dysphagia and difficulty swallowing present major obstacles to oral drug administration in geriatric and pediatric patients. This is especially challenging given the increased morbidity and need for medication with advanced age: an estimated 65% patients over 65 years of age are taking at least two medications, with 37% taking more than five^49, 50^. Difficulty taking pills may drive patients to skip doses or modify them in ways that dangerously alter drug pharmacokinetics. In this work, we developed a drinkable liquid formulation system, called LIFT hydrogels, capable of transitioning from liquid-to-solid upon mixing with ingested crosslinkers in the stomach. LIFT hydrogels have the advantages of solid formulations which confer enhanced gastric retention, protection against gastrointestinal proteases, toughness, capacity to control drug release, as well as co-encapsulation of drug with excipients. To realize LIFT hydrogels, we used FDA-approved or GRAS materials: alginate and 4-arm PEG-maleimide as hydrogel networks, and calcium chloride and DMSA or PEG dithiol as crosslinkers. The alginate/PEG solution remains a liquid until contact with the crosslinker solution within the stomach, facilitating a transition from a liquid to a tough hydrogel.

The gastric environment exhibits some features amenable for *in situ* crosslinking reactions. The stomach is temperature-controlled at 37 °C, which can accelerate maleimide/thiol thioether formation^51, 52^; the stomach is also mechanically active and its movement could facilitate mixing of the two ingested solutions^17^. We demonstrate that crosslinking of both alginate and PEG networks readily occur in *ex vivo* porcine gastric fluid and *in vivo* in porcine stomachs, which underscore the robustness of the calcium- and dithiol-mediated crosslinking reaction of the alginate and 4-arm PEG-maleimide networks. While LIFT hydrogel crosslinking and mechanical properties were dependent on the proportion of gastric fluid volume, this may be diluted through greater volumes of crosslinker. The fasted stomach contains 25-35 mL of gastric fluid^26, 53^, which after ingestion of a 200 mL crosslinker solution is diluted to 11-15%. This crosslinker volume is less than the volume of a typical drink can (355 mL), and this proportion of gastric fluid is well within the range capable of crosslinking both networks. Notably, these reactions do not generate side products, and the hydrogels did not appear to be toxic to cultured gastrointestinal epithelial, kidney, and liver cells, nor cause clinical or laboratory signals in pigs up to 48 h after administration.

Gastric drug depots should be able to withstand compressive forces within the stomach to preserve depot integrity. While hydrogels with dual polymer networks are mechanically stronger than single-network hydrogels^54, 55^, current strategies to formulate dual network hydrogels from orally administered liquids have not yet been described. Liquid systems have generally relied on single-network hydrogels of alginate, gellan, and karaya gum that are crosslinked by calcium.^14–16, 56^ In an alternative strategy, Li *et al.* utilized pH-triggered unmasking of multivalent cyclodextrin to undergo gelation with multivalent adamantane in acidic conditions^57^; however, the liberated masking group will need to be characterized for safety before application. While we and others have developed orally administrable tough hydrogels^55, 58, 59^, these require templated radical polymerization of toxic acrylamide monomer that cannot be safely performed *in vivo* and are dosed as a solid^60^. Other hydrogel systems have been designed that require UV light to facilitate polymer crosslinking^61, 62^, utilize polyacrylamide as a polymer network^37^, require a specific construction of hydrogel components^59^, or are enzymatically polymerized^63, 64^. While these hydrogels are mechanically tough, they either require a pre-solidified dosage format or are challenging and unsafe to crosslink and gel *in situ*. This work bridges this gap and enables liquid formulation of a tough hydrogel system. Assuming a spherical hydrogel, we calculate that reported gastric stresses (~13 kPa) would cause a strain of 5-10% in LIFT hydrogels, which should not permanently deform these hydrogels^17^.

While nano- and microparticle liquid suspensions have been developed, an important feature that this system confers is the *in situ* gelation of macroscale structures, which are important to minimize exposure of the dosage forms to gastric fluid. Given the same volume of material, nano- and microscale dosage forms result in significantly greater surface area-to-volume ratios compared to macroscale forms^65^. Economou *et al.* showed that the dissolution rate of CaCO_3_ particles was size-dependent, with larger 2-4 mm particles dissolving slower than <250 μm particles in acid^66^. Therefore, *in situ* gelation of macrostructures could be advantageous and enable protection of encapsulated therapeutics through size and geometry. Moreover, the formation of macroscale solids could prolong the gastric retention of encapsulated drugs compared to nano- and microparticulate liquid suspensions^19^.

This work can alter oral small molecule pharmacokinetics and prolong the function of biological drugs within the stomach. Patients who have difficulty swallowing solids may resort to crushing their pills, which results in dramatically altered pharmacokinetics that may cause severe complications and death^9, 11, 12^. Here we show that LIFT hydrogels modify pharmacokinetics by delaying and reducing the maximum drug plasma concentration while achieving a comparable AUC drug dose as free drug. This is significant because high drug concentrations can result in adverse side effects and impact treatment tolerability^67, 68^, and is applicable for drugs in which efficacy is driven by AUC and not blood concentration (e.g., tetracyclines, cytotoxics)^69, 70^ or drugs that require controlled release to stay within a safe therapeutic window. Given that a range of molecules are in equilibrium between the blood and gastrointestinal tract^71^, systems that modulate these molecules within the stomach could significantly impact healthcare via a noninvasive route. Indeed, engineered bacteria can sense blood within the stomach, remove nitrogenous waste, or metabolize excess phenylalanine^42, 45, 72, 73^. Oral enzyme and bacterial therapies are being developed for the treatment of hyperoxaluria and phenylketonuria, and are also used to treat patients with exocrine pancreatic insufficiency^74–76^. Coupling LIFT hydrogels with these therapeutics could alter drug pharmacokinetics and prolong both their residence and function within the gastrointestinal tract in a tough form factor. In doing so, LIFT hydrogels could expand the accessibility of these therapeutics to patients who otherwise have difficulty swallowing solids.

The chemistry of the LIFT hydrogels is robust, flexible, and tailorable. Here, we establish DMSA as a novel and FDA-approved small molecule crosslinker for these hydrogels, as well as a PEG-dithiol. Both crosslinkers were able to crosslink the 4-arm PEG-maleimide when mixed within the gastric cavity. The inclusion of PEG also enables facile covalent conjugation of drugs and other modulators using commercially available, functionalized multi-arm PEGs while still acting as a crosslinker. While maleimide-thiol reactions are rapid^77^, inclusion of the alginate network not only augments the mechanical properties of the hydrogel, but also its immediate formation in calcium solution acts as a “template” that facilitates retention and crosslinking of the slower-forming maleimide-thiol thioether bond. PEG networks typically require highly defined maleimide:thiol ratios for efficient gelation that is challenging to implement in an oral setting^52, 77^; here, templating within alginate likely enables gradual diffusion of the dithiol crosslinkers into the hydrogel and subsequent formation of the PEG network.

By overcoming the “ship-in-a-bottle” problem, LIFT hydrogels could expand access to medications for patients who have difficulty swallowing and bridge the advantages of solid and liquid drug formulations. Through careful selection of materials, LIFT hydrogels comprise two biocompatible polymer networks that are able to crosslink *in situ* within the stomach, resulting in a strong hydrogel that can facilitate localization of drugs and excipients and withstand the compressive forces of the gastrointestinal tract. LIFT hydrogels are safe, and are capable of modulating small molecule release and protecting therapeutic enzymes and bacteria in the stomach of large animals. We envision that LIFT hydrogels and their flexible chemistries may be a useful strategy with applications in gastric drug modulation and delivery, weight loss, and protection of encapsulated biologics.

## ACKNOWLEDGEMENTS

This work was funded in part by the Bill & Melinda Gates Foundation Grant INV-009529. M.J. was supported by the Translational Research Institute of Space Health through Cooperative Agreement NNX16AO69A. We would like to thank Ameya Kirtane for insightful discussions.

## Author contributions

G.W.L. and M.J.P. conceived and designed the hydrogel. G.W.L. and M.J.P. performed experiments. V.R.F. performed rheology experiments. M.J. made the bioluminescent bacterial strain and helped design and analyze the bacterial protection experiments. J.L.P.K., K.I., W.A.M.M., G.N.W., J.J., N.Fabian., and A.H. assisted with *in vivo* experimentation, A.L., J.M., and N.Fitzgerald assisted with bioanalytics, and J.H.C and C.K.S. performed cell culture experiments. R.S.L. and G.T. provided funding and supervised. All authors contributed to and reviewed the manuscript.

## CONCISE MATERIALS AND METHODS

### Chemicals

Poly(ethylene glycol) (PEG)-dithiol (1 kDa) was purchased from Biopharma PEG, 4-arm PEG-maleimide (20 kDa) was purchased from JenKem Technology USA, Laysan Bio, Inc, and Creative PEGWorks, and alginate (71238), trypsin (T7409), and β-galactosidase (G5160) were purchased from MilliporeSigma. Alginate solutions were prepared in ddH_2_O by vigorous heating and stirring. Calcium carbonate and dimercaptosuccinic acid (DMSA) were purchased from ACROS Organics, and o-nitrophenyl β-D-galactopyranoside (ONPG) was purchased from Cayman Chemical. Lumefantrine was purchased from Fisher Scientific, and halofantrine was purchased from MedChemExpress.

### *In vitro* LIFT hydrogel formation and characterization

A polymer solution of 0.5% w/v alginate and 4-arm PEG-maleimide (0-10% w/v) was prepared in ddH_2_O; to form hydrogels, 60 μL of this solution was cast into 1 mL of crosslinker solution (200 mM CaCl_2_, 10 mM PEG-dithiol or DMSA) using a pipette and incubated for 10-20 min at 37 °C, 50 RPM. After, the resulting hydrogels were washed with ddH_2_O. In some experiments, hydrogels were cast in 0-100% v/v of porcine gastric fluid diluted with ddH_2_O. Concentrated stocks of crosslinker were added to porcine gastric fluid to achieve the stated crosslinker concentrations and % v/v gastric fluid.

Mechanical compression tests were performed using an Instron instrument as previously described with modifications.^78^ The gauge length was determined with a digital caliper, and displacement was applied at a rate of 0.05 mm/s until 90% strain. Cyclic compression measurement was performed with a displacement rate of 0.05 mm/s and 5 cycles of 90% strain.

To facilitate rheological characterization, pre-crosslinked LIFT hydrogels were prepared by casting a solution of 0.5% alginate/5% w/v 4-arm PEG-maleimide with 15 mM CaCO_3_/30 mM glucono-δ-lactone into a 100-mm Petri dish. After 1 h incubation at room temperature, hydrogel samples were made using an 8-mm diameter biopsy punch. Oscillatory rheology studies were performed with a Discovery Series Hybrid Rheometer from TA Instruments. Samples were measured using 8-mm parallel plates fully submerged in a 5 mL bath of crosslinker solution (200 mM CaCl2/10 mM PEG-dithiol) at 37 °C. We reasoned that the 8-mm parallel plates (smallest available size) would minimize unexposed surface area at the top and bottom faces of the sample, and therefore best represent crosslinking dynamics *in vivo*. Data was collected for 1 h with a frequency of 10 rad/s and strain of 1%.

### Model encapsulation and release

The following model encapsulants were mixed at a 10 mg/mL concentration in either alginate or LIFT polymer solutions: 155-kDa tetramethylrhodamine isothiocyanate-dextran (TRITC-dextran, MilliporeSigma), and 20- and 100-nm fluorescent carboxylated polystyrene nanoparticles (ThermoFisher). Hydrogels were formed as described above, transferred to simulated gastric fluid (SGF: 34 mM NaCl pH 1.77) or simulated intestinal fluid (SIF, Cole-Parmer), and then incubated at 37 °C, 50 RPM. The supernatant was sampled at various timepoints with replacement. For each hydrogel and model encapsulant, three separate experiments were performed simultaneously, and release was calculated according to respective standards diluted in either SGF or SIF.

### Cytotoxicity

Cytotoxicity of gel constituents (4-arm PEG-maleimide, PEG-dithiol, DMSA, CaCl_2_) was determined for 4 different cell lines: Caco-2, HT-29, Hepa1-6, and CV-1. Cell lines were mycoplasma-tested as negative prior to use, and genomically fingerprinted, where possible, to verify their identity. Alginate was unable to be tested due to its viscosity and incompatibility with robotic fluid handlers. Cells were plated at 15,000 cells/well in DMEM+10% FBS using robotic handlers (Tecan Evo 150) and incubated overnight. After, cells were incubated in the indicated treatments and concentrations for 24 h in DMEM+10% FBS, and viability was quantified using CellTiter-Glo (Promega), which uses intracellular ATP levels as a surrogate for viability, and a plate reader (Tecan Infinite Pro 1000). Viability was calculated as a percentage of untreated cells.

### *In vivo* experimentation

All animal studies were performed only after MIT CAC review and approval and under veterinary supervision. Specific methods and treatments for characterization, lumefantrine, lactase, and bacteria studies in rats and pigs are described within their respective sections. Female Yorkshire pigs 50-100 kg and rats >400 g were used.

### *In vivo* LIFT hydrogel formation and characterization

Hydrogels were administered into stomachs of anesthetized pigs via endoscopy. To facilitate visualization, gastric fluid was removed. Crosslinker solution (200 mL, 200 mM CaCl_2_ and 10 mM DMSA or PEG-dithiol) was first administered, and then the endoscope was purged with air and water. After, 20-40 mL of hydrogel solution (typically 0.5% alginate/5% w/v 4-arm PEG-maleimide) was similarly administered. In some experiments, the order was reversed. For mechanical characterization, pigs were sacrificed 6-8 h after hydrogel administration, and the hydrogels were retrieved and tested as described above.

To monitor hydrogel retention kinetics in the porcine gastrointestinal tract and for acute toxicity, hydrogels were loaded with barium sulfate (20% w/v) for X-ray imaging, and images were collected immediately after administration, 4-5 h, and on days 1 and 2. Serum was collected before hydrogel administration (baseline) and on days 1 and 2 for metabolic analysis. Throughout, pigs were clinically monitored for gastrointestinal symptoms (e.g., inappetence, vomiting).

### Encapsulated lumefantrine pharmacokinetics

Pigs were dosed under anesthesia via endoscopy with the following treatments: free lumefantrine, lumefantrine encapsulated in alginate hydrogels, and lumefantrine encapsulated in LIFT hydrogels (*n* = 3 each). All pigs were dosed with a total of 960 mg lumefantrine. For free lumefantrine, drug powder was weighed and placed across three gelatin capsules. For hydrogel formulations, lumefantrine powder was suspended in polymer solution (0.5% w/v alginate or 0.5% alginate/5% w/v 4-arm PEG-maleimide), mixed, and administered after crosslinker solution (200 mM CaCl_2_/10 mM DMSA). Blood was sampled from a central jugular catheter at the indicated time points, and lumefantrine area under the curve was calculated by the trapezoidal rule.

Plasma lumefantrine was separated via high pressure liquid chromatography (HPLC) and quantified with an Agilent 6495A triple quadrupole mass spectrometer equipped with a sheath gas electrospray ionization (AJS-ESI) (Agilent Technologies, Santa Clara, CA). Samples were injected at a 5 μL injection volume. Chromatography was performed on an Acquity BEH C18 column (2.1×50 mm, dp 1.8 μm, Waters, Milford, MA), heated to 50 °C, with a binary mobile phase composed of 0.1% formic acid in water (v/v, A) and 5% tetrahydrofuran in methanol (v/v, B). The mobile phase was pumped at 0.5 mL/min and gradient programmed as: 0 min, 5% B; 5 min, 95% B. The total method runtime was 7 min with a 2 min re-equilibration time between injections. For positive ionization ESI source conditions, the iFunnel high pressure radiofrequency was set to 150 V, and low pressure set to 60 V. Nebulizer drying gas temperature was set to 210 °C with a flow rate of 15 L/min at 35 psig. Sheath gas temperature was set to 250 °C with a flow rate of 12 L/min. Nozzle voltage was set to 1500 V and capillary voltage was set to 3500 V. Dynamic multiple reaction-monitoring was used to quantify analytes, using nitrogen as the collision gas. Lumefantrine was quantified at transitions 528.16 to 510.00 m/z at 28 collision energy (CE), with a qualifier transition from 528.16 to 383.00 m/z (40 CE). Halofantrine was used as an internal standard and quantified with the 500.18 to 142.10 m/z transition (24 CE) and qualified with the 500.18 to 100.10 m/z transition. All transitions used a cell accelerator voltage of 4. Data analysis was performed with MassHunter B10.1 (Agilent Technologies, Santa Clara, CA). Linear calibration curves were weighted by the reciprocal of the standard concentrations used, i.e., 1/x.

A ten-point calibration curve of halofantrine and lumefantrine was prepared with concentrations ranging from 1 to 2500 ng/mL. For plasma sample preparation, 250 μL of plasma, 20 μL of halofantrine at 2500 ng/mL and 730 μL of 90:10 methanol:tetrahydrofuran was added for protein precipitation. Samples were vortexed and centrifuged at 15,000×g for 15 min. The resulting supernatant (200 μL) was transferred to glass vials for analysis.

### Lactase activity after dithiol molecule treatment

Lactase (18 μg, 60 *μ*L) was added to DMSA or PEG-dithiol to a final DMSA/PEG-dithiol concentration of 2.5, 5, or 10 mM. Treatments were incubated at 37 °C for 20 min. Lactase activity was assayed by adding 60 μL of 5 mM ONPG and incubation for 1 min at room temperature. After, 300 μL of 1 M Na_2_CO_3_ was added to stop the reaction, and the absorbance of the solution was read at λ = 420 nm.

### Lactase encapsulation in LIFT hydrogels

The effect of acid on enzyme activity was determined by treating lactase (0.24 mg, 60 μL) with either SGF or PBS and incubating at 37 °C, 50 RPM for various times. Enzymatic activity after incubation was determined by adding 60 μL of 5 mM ONPG and incubation for 1 min at room temperature. After, 300 μL of 1 M Na_2_CO_3_ was added to stop the reaction, and the absorbance of the solution was read at λ= 420 nm. For *in vitro* hydrogel experiments, lactase (0.20 mg) was suspended in 60 μL hydrogel solution (0.5% alginate/5% w/v 4-arm PEG-maleimide) and cast in crosslinker solution (200 mM CaCl_2_/10 mM PEG-SH). Alginate-only hydrogels were prepared in 200 mM CaCl_2_ solution only, and both hydrogels were prepared with and without CaCO_3_ (42.68 mg/mL). Hydrogels were then challenged with SGF for 1 h at 37 °C. After acid incubation, enzymatic activity was quantified as above. For trypsin challenge experiments, lactase-containing hydrogels (60 μL, 0.20 mg lactase) were prepared and incubated with trypsin (40 mg/mL) for 6 h at 37 °C. Free lactase and alginate-only hydrogels were included as controls. Lactase enzyme activity was quantified as previously described, and compared between trypsin-treated samples and naive samples to determine relative absorbance.

Encapsulated enzyme activity was tested in rat and porcine models. Rats were fasted overnight prior to administration. The following day, 3 mL of crosslinker solution (200 mM CaCl_2_, 10 mM PEG-dithiol) was administered via oral gavage immediately followed by 1 mL of hydrogel solution (0.5% alginate/5% w/v 4-arm PEG-maleimide with 0.24 mg lactase). Calcium carbonate (42.69 mg) was included either in the crosslinker solution (separate) or in the hydrogel solution (co-encapsulated). After 1 or 2 h, rats were euthanized, and the hydrogels were collected. Hydrogels were weighed and minced, and enzymatic activity was quantified as described above and normalized by hydrogel mass. Encapsulated enzyme activity was also tested in Yorkshire pigs. Hydrogels were administered via endoscopy into the stomach: first, 200 mL of crosslinker (200 mM CaCl_2_/10 mM PEG-dithiol) was administered followed by 20 mL of 0.5% alginate/5% w/v 4-arm PEG-maleimide containing lactase (40.45 mg) with or without co-encapsulated CaCO_3_ (2 g). After 6 h, hydrogels were retrieved, and lactase activity was quantified as described above.

### Bacteria encapsulation in LIFT hydrogels

*E. coli* Nissle 1917 was isolated from commercially available Mutaflor capsules on LB-agar plates (BD 240230). This strain was transformed with the plasmid pAKlux2 (Addgene 14080) to create a constitutively bioluminescent *E. coli* Nissle 1917 strain. Bioluminescent *E. coli* Nissle 1917 was routinely cultured on LB-agar plates at 37 °C or in LB in culture tubes shaken at 37 °C containing ampicillin (100 μg/mL). The bacterial concentration in overnight cultures was determined by measuring the OD_600_, and the cells were pelleted by centrifugation and resuspended in PBS at the target concentration.

To determine bacteria activity in SGF pH 1.77, bacteria (1×10^8^ CFU, 68 μL) was added to 132 μL SGF or PBS and incubated for the indicated timepoints. Bioluminescence was recorded using a plate reader (Infinite 200, Tecan).

For *in vitro* hydrogel experiments, bacteria (1×10^8^ CFU) was suspended in 60 μL hydrogel solution (0.5% alginate/5% w/v 4-arm PEG-maleimide) and cast in 1 mL of crosslinker solution (200 mM CaCl_2_/10 mM PEG-SH) with and without CaCO_3_ (7.5 mg/mL). Hydrogels were transferred to 100 μL SGF for 3 h at 37 °C, 100 RPM. After acid incubation, hydrogels were transferred to a white 96-well plate and analyzed for bioluminescence. Hydrogels were then transferred to 1 mL LB media and incubated for 4 h, 37 °C, 100 RPM. After, the supernatant was collected and analyzed for bioluminescence.

Encapsulated bacteria activity was tested in porcine models. Hydrogels were administered into the stomach of anesthetized pigs via endoscopy: first, 200 mL of crosslinker (200 mM CaCl_2_/10 mM PEG-dithiol) was administered followed by 20 mL of 0.5% alginate/5% w/v 4-arm PEG-maleimide containing bacteria (1.6×10^10^ CFU) with or without co-encapsulated CaCO_3_ (2 g). After 6-8 h, hydrogels were retrieved, and bacteria viability was quantified using an *in vivo* imaging system (PerkinElmer). This experiment was repeated 3 times in different pigs.

### Statistical analysis

For all tests, an α = 0.05 was set for statistical significance. Single comparison tests were performed using a Student’s *t*-test, and multiple comparisons were performed using a one-way ANOVA with post-hoc Tukey’s multiple comparisons test.

## REFERENCES

1. Liu, G., Franssen, E., Fitch, M.I. & Warner, E. Patient preferences for oral versus intravenous palliative chemotherapy. J Clin Oncol 15, 110–115 (1997).

2. Eek, D. et al. Patient-reported preferences for oral versus intravenous administration for the treatment of cancer: a review of the literature. Patient Prefer Adherence 10, 1609–1621 (2016).

3. Alyami, H. et al. Dosage form preference consultation study in children and young adults: paving the way for patient-centred and patient-informed dosage form development. Eur J Hosp Pharm 24, 332–337 (2017).

4. Rubin, R.R., Peyrot, M., Kruger, D.F. & Travis, L.B. Barriers to insulin injection therapy: patient and health care provider perspectives. Diabetes Educ 35, 1014–1022 (2009).

5. Fu, A.Z., Qiu, Y. & Radican, L. Impact of fear of insulin or fear of injection on treatment outcomes of patients with diabetes. Curr Med Res Opin 25, 1413–1420 (2009).

6. Zhong, W. et al. Age and sex patterns of drug prescribing in a defined American population. Mayo Clin Proc 88, 697–707 (2013).

7. Patel, A., Jacobsen, L., Jhaveri, R. & Bradford, K.K. Effectiveness of pediatric pill swallowing interventions: a systematic review. Pediatrics 135, 883–889 (2015).

8. Alessandrini, E. et al. Children’s Preferences for Oral Dosage Forms and Their Involvement in Formulation Research via EPTRI (European Paediatric Translational Research Infrastructure). Pharmaceutics 13, 730–730 (2021).

9. Schiele, J.T., Quinzler, R., Klimm, H.D., Pruszydlo, M.G. & Haefeli, W.E. Difficulties swallowing solid oral dosage forms in a general practice population: prevalence, causes, and relationship to dosage forms. Eur J Clin Pharmacol 69, 937–948 (2013).

10. Adkins, C. et al. Prevalence and Characteristics of Dysphagia Based on a Population-Based Survey. Clin Gastroenterol Hepatol 18, 1970–1979 e1972 (2020).

11. Cleary, J.D., Evans, P.C., Hikal, A.H. & Chapman, S.W. Administration of crushed extended-release pentoxifylline tablets: bioavailability and adverse effects. Am J Health Syst Pharm 56, 1529–1534 (1999).

12. Logrippo, S. et al. Oral drug therapy in elderly with dysphagia: between a rock and a hard place! Clin Interv Aging 12, 241–251 (2017).

13. Lajoinie, A., Henin, E., Kassai, B. & Terry, D. Solid oral forms availability in children: a cost saving investigation. Br J Clin Pharmacol 78, 1080–1089 (2014).

14. Xu, X. et al. Intragastric amorphous calcium carbonate consumption triggered generation of in situ hydrogel piece for sustained drug release. Int J Pharm 590, 119880 (2020).

15. Kubo, W., Miyazaki, S. & Attwood, D. Oral sustained delivery of paracetamol from in situ-gelling gellan and sodium alginate formulations. Int J Pharm 258, 55–64 (2003).

16. Foster, K.A. et al. Utility of in situ sodium alginate/karaya gum gels to facilitate gastric retention in rodents. Int J Pharm 434, 406–412 (2012).

17. Houghton, L.A. et al. Motor activity of the gastric antrum, pylorus, and duodenum under fasted conditions and after a liquid meal. Gastroenterology 94, 1276–1284 (1988).

18. Hong, S. et al. 3D Printing of Highly Stretchable and Tough Hydrogels into Complex, Cellularized Structures. Adv Mater 27, 4035–4040 (2015).

19. Bennink, R. et al. Comparison of total and compartmental gastric emptying and antral motility between healthy men and women. Eur J Nucl Med 25, 1293–1299 (1998).

20. Martinsen, T.C., Fossmark, R. & Waldum, H.L. The Phylogeny and Biological Function of Gastric Juice-Microbiological Consequences of Removing Gastric Acid. Int J Mol Sci 20(2019).

21. Lee, K.Y. & Mooney, D.J. Alginate: properties and biomedical applications. Prog Polym Sci 37, 106–126 (2012).

22. McGraw, T. Safety of polyethylene glycol 3350 solution in chronic constipation: randomized, placebo-controlled trial. Clin Exp Gastroenterol 9, 173–180 (2016).

23. Stephan, M.T., Moon, J.J., Um, S.H., Bershteyn, A. & Irvine, D.J. Therapeutic cell engineering with surface-conjugated synthetic nanoparticles. Nature Medicine 16, 1035–1041 (2010).

24. Chisolm, J.J., Jr. Safety and efficacy of meso-2,3-dimercaptosuccinic acid (DMSA) in children with elevated blood lead concentrations. J Toxicol Clin Toxicol 38, 365–375 (2000).

25. Lifshitz, M., Hashkanazi, R. & Phillip, M. The effect of 2,3 dimercaptosuccinic acid in the treatment of lead poisoning in adults. Ann Med 29, 83–85 (1997).

26. Mudie, D.M. et al. Quantification of gastrointestinal liquid volumes and distribution following a 240 mL dose of water in the fasted state. Mol Pharm 11, 3039–3047 (2014).

27. Braim, S. et al. Lactoferrin-Loaded Alginate Microparticles to Target Clostridioides difficile Infection. J Pharm Sci 108, 2438–2446 (2019).

28. Ilgin, P., Ozay, H. & Ozay, O. Synthesis and characterization of pH responsive alginate based-hydrogels as oral drug delivery carrier. Journal of Polymer Research 27, 251 (2020).

29. Ziegler, A., Gonzalez, L. & Blikslager, A. Large Animal Models: The Key to Translational Discovery in Digestive Disease Research. Cellular and Molecular Gastroenterology and Hepatology 2, 716–724 (2016).

30. Davis, S.S., Illum, L. & Hinchcliffe, M. Gastrointestinal transit of dosage forms in the pig. J Pharm Pharmacol 53, 33–39 (2001).

31. Kadiyala, I. & Tan, E. Formulation approaches in mitigating toxicity of orally administrated drugs. Pharm Dev Technol 18, 305–312 (2013).

32. Vllasaliu, D., Thanou, M., Stolnik, S. & Fowler, R. Recent advances in oral delivery of biologics: nanomedicine and physical modes of delivery. Expert Opin Drug Deliv 15, 759–770 (2018).

33. Lomer, M.C., Parkes, G.C. & Sanderson, J.D. Review article: lactose intolerance in clinical practice--myths and realities. Aliment Pharmacol Ther 27, 93–103 (2008).

34. Perissinato, A.G., Morais Ruela, A.L., Pereira, G.R., Garcia, J.S. & Trevisan, M.G. Simple Strategy to Protect Lactase Activity in Solid Formulation. Curr Drug Deliv 15, 215–218 (2018).

35. Nichele, V., Signoretto, M. & Ghedini, E. β-Galactosidase entrapment in silica gel matrices for a more effective treatment of lactose intolerance. Journal of Molecular Catalysis B: Enzymatic 71, 10–15 (2011).

36. Zhang, Z., Zhang, R. & McClements, D.J. Lactase (β-galactosidase) encapsulation in hydrogel beads with controlled internal pH microenvironments: Impact of bead characteristics on enzyme activity. Food Hydrocolloids 67, 85–93 (2017).

37. Tang, T.-C. et al. Hydrogel-based biocontainment of bacteria for continuous sensing and computation. Nature Chemical Biology 17, 724–731 (2021).

38. Hu, Y. et al. A double-layer hydrogel based on alginate-carboxymethyl cellulose and synthetic polymer as sustained drug delivery system. Sci Rep 11, 9142 (2021).

39. Mahou, R. et al. Combined Electrostatic and Covalent Polymer Networks for Cell Microencapsulation. Macromolecular Symposia 329, 49–57 (2013).

40. Fritz, K., Taylor, K. & Parmar, M. in StatPearls (Treasure Island (FL); 2022).

41. Anselmo, A.C., McHugh, K.J., Webster, J., Langer, R. & Jaklenec, A. Layer-by-Layer Encapsulation of Probiotics for Delivery to the Microbiome. Adv Mater 28, 9486–9490 (2016).

42. Kurtz, C.B. et al. An engineered E. coli Nissle improves hyperammonemia and survival in mice and shows dose-dependent exposure in healthy humans. Sci Transl Med 11 (2019).

43. Adolfsen, K.J. et al. Improvement of a synthetic live bacterial therapeutic for phenylketonuria with biosensor-enabled enzyme engineering. Nature Communications 12, 6215 (2021).

44. Canale, F.P. et al. Metabolic modulation of tumours with engineered bacteria for immunotherapy. Nature 598, 662–666 (2021).

45. Puurunen, M.K. et al. Safety and pharmacodynamics of an engineered E. coli Nissle for the treatment of phenylketonuria: a first-in-human phase 1/2a study. Nat Metab 3, 1125–1132 (2021).

46. Merchant, H.A. et al. Assessment of gastrointestinal pH, fluid and lymphoid tissue in the guinea pig, rabbit and pig, and implications for their use in drug development. Eur J Pharm Sci 42, 3–10 (2011).

47. Yang, S.Y. et al. Powering Implantable and Ingestible Electronics. Adv Funct Mater 31 (2021).

48. Gardner, J.D., Ciociola, A.A. & Robinson, M. Measurement of meal-stimulated gastric acid secretion by in vivo gastric autotitration. J Appl Physiol (1985) 92, 427–434 (2002).

49. Young, E.H., Pan, S., Yap, A.G., Reveles, K.R. & Bhakta, K. Polypharmacy prevalence in older adults seen in United States physician offices from 2009 to 2016. PLoS One 16, e0255642 (2021).

50. Maher, R.L., Hanlon, J. & Hajjar, E.R. Clinical consequences of polypharmacy in elderly. Expert Opin Drug Saf 13, 57–65 (2014).

51. Babaee, S. et al. Temperature-responsive biometamaterials for gastrointestinal applications. Sci Transl Med 11 (2019).

52. Darling, N.J., Hung, Y.S., Sharma, S. & Segura, T. Controlling the kinetics of thiol-maleimide Michael-type addition gelation kinetics for the generation of homogenous poly(ethylene glycol) hydrogels. Biomaterials 101, 199–206 (2016).

53. Grimm, M., Koziolek, M., Kühn, J.P. & Weitschies, W. Interindividual and intraindividual variability of fasted state gastric fluid volume and gastric emptying of water. Eur J Pharm Biopharm 127, 309–317 (2018).

54. Macdougall, L.J. et al. Self-healing, stretchable and robust interpenetrating network hydrogels. Biomater Sci 6, 2932–2937 (2018).

55. Liu, J. et al. Triggerable tough hydrogels for gastric resident dosage forms. Nature Communications 8, 124 (2017).

56. Katayama, H. et al. Sustained Release Liquid Preparation Using Sodium Alginate for Eradication of Helicobacter pyroli. Biological & Pharmaceutical Bulletin 22, 55–60 (1999).

57. Li, Z. et al. Hydrogel Transformed from Nanoparticles for Prevention of Tissue Injury and Treatment of Inflammatory Diseases. Adv Mater 34, e2109178 (2022).

58. Jin, X., Wei, C., Wu, C. & Zhang, W. Gastric fluid-induced double network hydrogel with high swelling ratio and long-term mechanical stability. Composites Part B: Engineering 236, 109816 (2022).

59. Liu, X. et al. Ingestible hydrogel device. Nat Commun 10, 493 (2019).

60. Besaratinia, A. & Pfeifer, G.P. A review of mechanisms of acrylamide carcinogenicity. Carcinogenesis 28, 519–528 (2007).

61. Mredha, M.T.I. et al. Anisotropic tough double network hydrogel from fish collagen and its spontaneous in vivo bonding to bone. Biomaterials 132, 85–95 (2017).

62. Aldana, A.A. et al. Biomimetic double network hydrogels: Combining dynamic and static crosslinks to enable biofabrication and control cell-matrix interactions. Journal of Polymer Science 59, 2832–2843 (2021).

63. Wei, Q. et al. One-pot preparation of double network hydrogels via enzyme-mediated polymerization and post-self-assembly for wound healing. J Mater Chem B 7, 6195–6201 (2019).

64. Geng, J. et al. A Fluorescent Biofunctional DNA Hydrogel Prepared by Enzymatic Polymerization. Adv Healthc Mater 7 (2018).

65. Laroui, H. et al. Nanomedicine in GI. Am J Physiol Gastrointest Liver Physiol 300, G371–383 (2011).

66. Economou, E.D., Evmiridis, N.P. & Vlessidis, A.G. Dissolution Kinetics of CaCO3 in Powder Form and Influence of Particle Size and Pretreatment on the Course of Dissolution. Industrial & Engineering Chemistry Research 35, 465–474 (1996).

67. Gatti, G. et al. The relationship between ritonavir plasma levels and side-effects: implications for therapeutic drug monitoring. AIDS 13 (1999).

68. Saetre, E., Perucca, E., Isojarvi, J., Gjerstad, L. & Group, L.A.M.S. An international multicenter randomized double-blind controlled trial of lamotrigine and sustained-release carbamazepine in the treatment of newly diagnosed epilepsy in the elderly. Epilepsia 48, 1292–1302 (2007).

69. Agwuh, K.N. & MacGowan, A. Pharmacokinetics and pharmacodynamics of the tetracyclines including glycylcyclines. J Antimicrob Chemother 58, 256–265 (2006).

70. Sadekar, S., Figueroa, I. & Tabrizi, M. Antibody Drug Conjugates: Application of Quantitative Pharmacology in Modality Design and Target Selection. AAPS J 17, 828–836 (2015).

71. Steiger, C. et al. Dynamic Monitoring of Systemic Biomarkers with Gastric Sensors. Adv Sci (Weinh) 8, e2102861 (2021).

72. Mimee, M. et al. An ingestible bacterial-electronic system to monitor gastrointestinal health. Science 360, 915–918 (2018).

73. Zheng, D.W. et al. An orally delivered microbial cocktail for the removal of nitrogenous metabolic waste in animal models of kidney failure. Nat Biomed Eng 4, 853–862 (2020).

74. Lingeman, J.E. et al. ALLN-177, oral enzyme therapy for hyperoxaluria. Int Urol Nephrol 51, 601–608 (2019).

75. Kim, W. et al. Trends in enzyme therapy for phenylketonuria. Mol Ther 10, 220–224 (2004).

76. Kuhn, R.J., Gelrud, A., Munck, A. & Caras, S. CREON (Pancrelipase Delayed-Release Capsules) for the treatment of exocrine pancreatic insufficiency. Adv Ther 27, 895–916 (2010).

77. Jansen, L.E., Negron-Pineiro, L.J., Galarza, S. & Peyton, S.R. Control of thiol-maleimide reaction kinetics in PEG hydrogel networks. Acta Biomater 70, 120–128 (2018).

78. Liu, J. et al. Triggerable tough hydrogels for gastric resident dosage forms. Nat Commun 8, 124 (2017).

