## Supplementary Information for "Drinkable, liquid *in situ*-forming and tough hydrogels for gastrointestinal therapeutics"

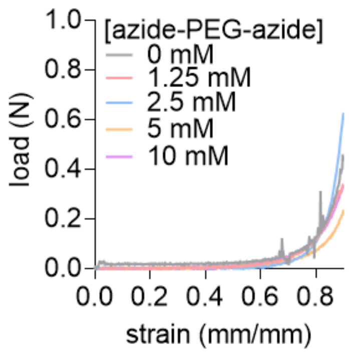


**Figure S1.** Load-strain curves of DBCO-azide PEG/alginate hydrogels. Hydrogels were formed by drop casting a solution of 0.5% alginate/5% w/v 4-arm PEG-DBCO into a solution of 200 mM $CaCl_2$ with the indicated concentrations of PEG diazide crosslinker. After casting, hydrogels were incubated at 37 °C for 20 min and then mechanically characterized by compression testing. Hydrogels crosslinked in the presence of PEG diazide were not mechanically different than the no PEG diazide control, indicating that the PEG network did not appreciably crosslink within 20 min. Moreover, all hydrogels remained permanently deformed (flattened) after 90% strain.

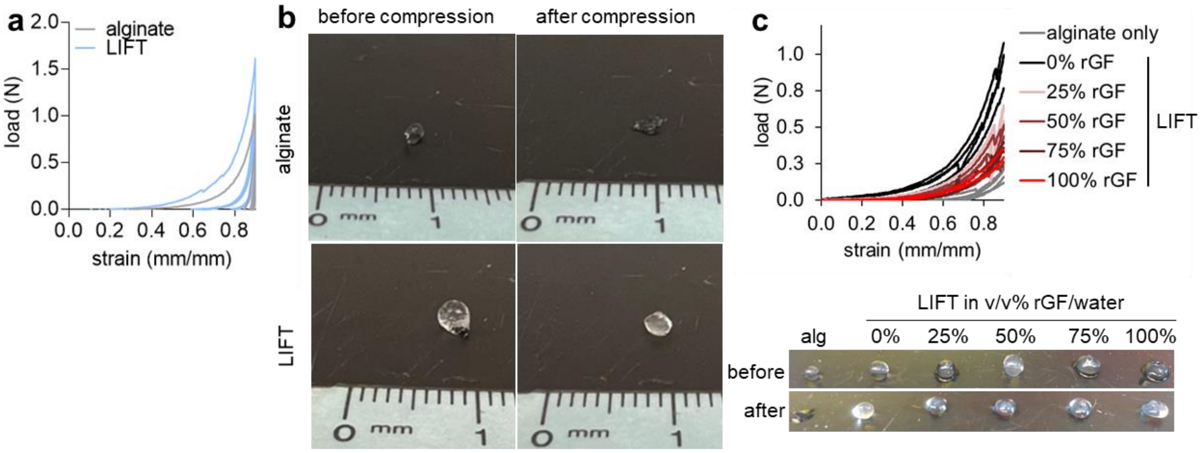


**Figure S2. A.** Load-strain curves of LIFT and alginate hydrogels throughout 5 cycles of 90% strain. Hydrogels were formed by drop casting of 0.5% alginate/5% w/v 4-arm PEG-maleimide into a solution of 200 mM $CaCl_2$/10 mM DMSA. After casting, hydrogels were incubated at 37 °C, 50 RPM, 20 min and then mechanically characterized by compression testing. While LIFT hydrogels exhibited some elasticity throughout multiple compressions, alginate hydrogels were permanently deformed after the first compression and were unable to sustain subsequent strains. **B.** Images of alginate and LIFT hydrogels before and after 90% cyclic strain. Whereas alginate hydrogels were permanently deformed, LIFT hydrogels exhibited some recovery. **C.** Top: load-strain curves of LIFT hydrogels after formation in real gastric fluid (rGF). Hydrogels were formed by drop casting of 0.5% alginate/5% w/v 4-arm PEG-maleimide into a solution of rGF with 100 mM $CaCl_2$/5 mM DMSA. To achieve the indicated rGF proportions, rGF was diluted with water. After casting, hydrogels were incubated at 37 °C, 50 RPM, 10 min and then mechanically characterized by compression testing. Despite formation in complicated gastric fluid, LIFT hydrogels formed in rGF were mechanically stronger than alginate hydrogels formed in 100 mM $CaCl_2$ without rGF. Bottom: hydrogels before and after 90% strain. While alginate hydrogels remained permanently flattened, LIFT hydrogels formed in rGF retained some degree of their original geometry.

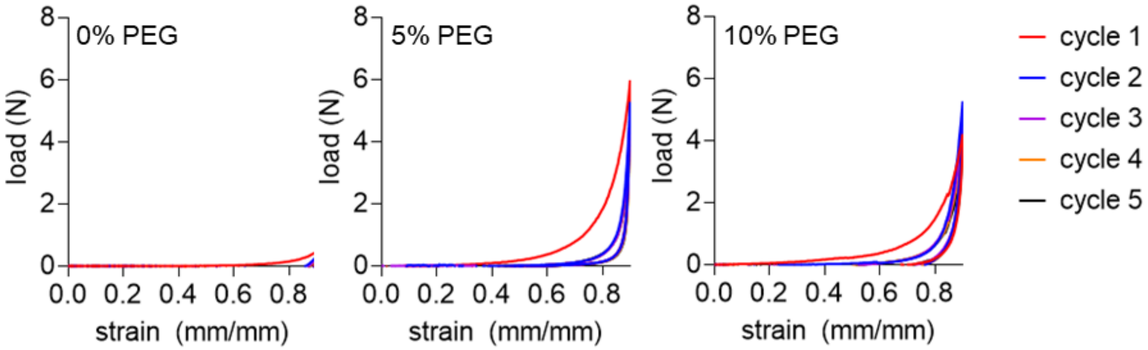


**Figure S3.** Load-strain curves of hydrogels throughout 5 cycles of 90% strain. Hydrogels were formed by drop casting of 0.5% w/v alginate and the indicated amount (% w/v) of 4-arm PEG-maleimide into a solution of 200 mM $CaCl_2$/10 mM PEG-dithiol. After casting, hydrogels were incubated at 37 °C, 50 RPM, 20 min and then mechanically characterized by compression testing. While hydrogels containing a crosslinked PEG network recovered a degree of their original geometry throughout multiple compressions, alginate hydrogels were permanently flattened after the first compression and were unable to sustain subsequent strains.

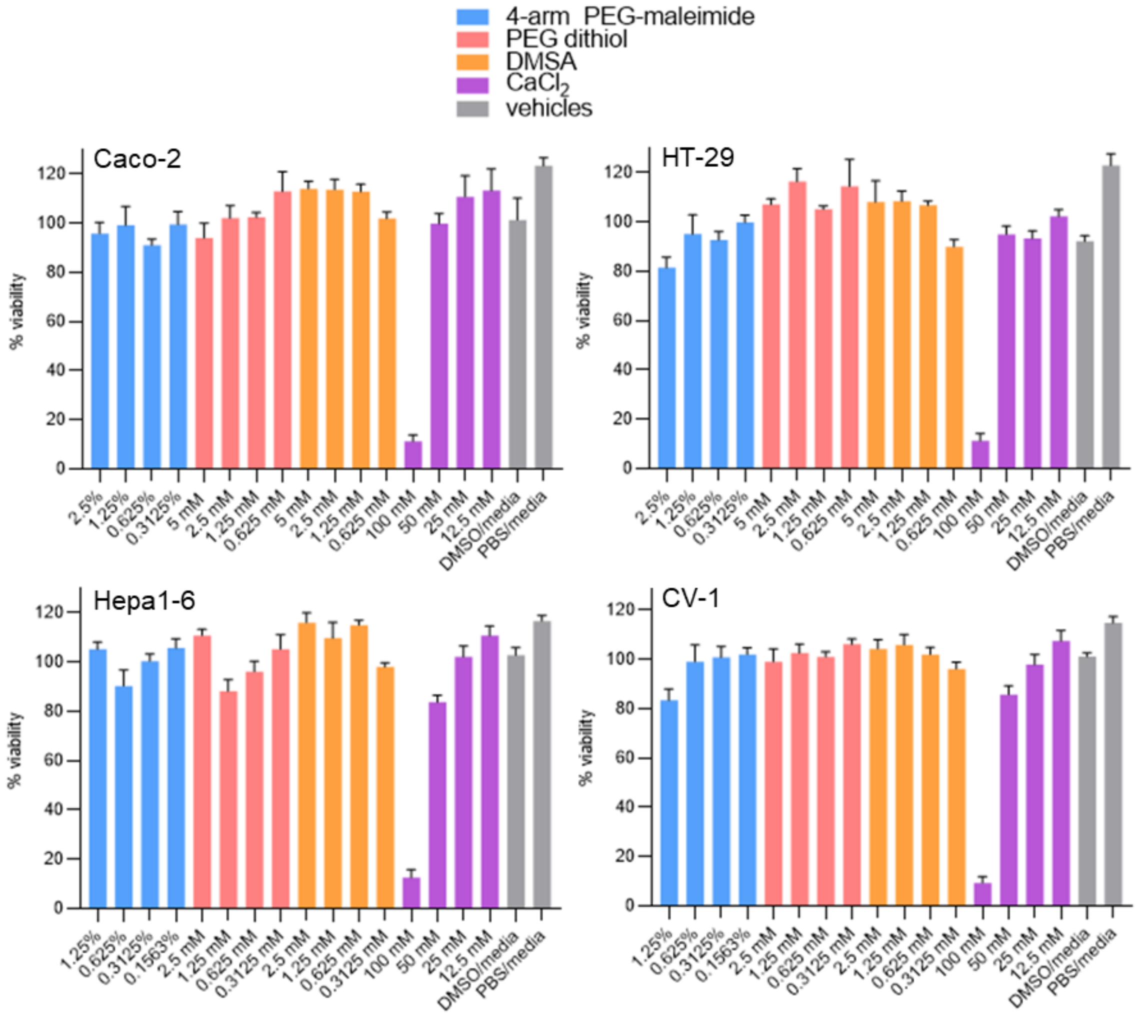


**Figure S4.** Viability of various cell lines after 24 h exposure to various concentrations of 4-arm PEG-maleimide, PEG dithiol, dimercaptosuccinic acid (DMSA), and $CaCl_2$. Cell lines tested were human colon epithelial Caco-2, human colon epithelial HT-29, mouse liver Hepa1-6, and monkey kidney CV-1 cells. An $n$ = 6 wells were tested for each treatment per cell line; viability was normalized to untreated cells.

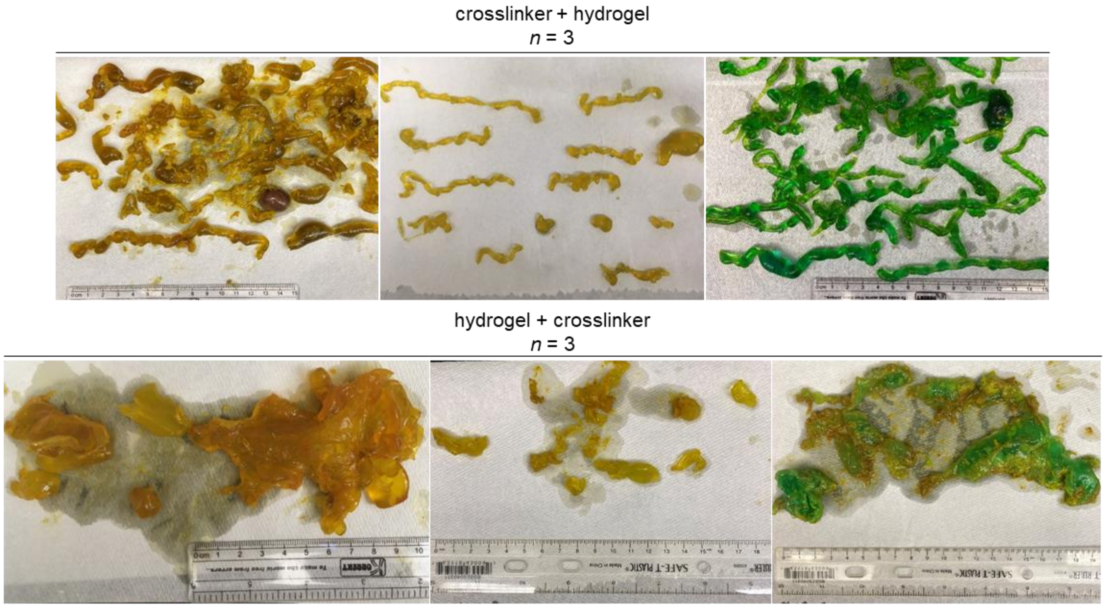


**Figure S5.** Structure study of LIFT hydrogels formed *in vivo*. Top: pigs ($n$ = 3) were administered crosslinker solution (200 mM $CaCl_2$/10 mM DMSA) followed by hydrogel solution (0.5% alginate/5% w/v 4-arm PEG-maleimide). In some experiments, green dye was added for color contrast. Bottom: pigs ($n$ = 3) were administered hydrogel solution followed by crosslinker solution. In some experiments, green dye was added for color contrast. Generally, the administration order crosslinker + hydrogel resulted in consistent “noodle-like” hydrogels, while hydrogel + crosslinker resulted in more heterogenous hydrogel shapes.

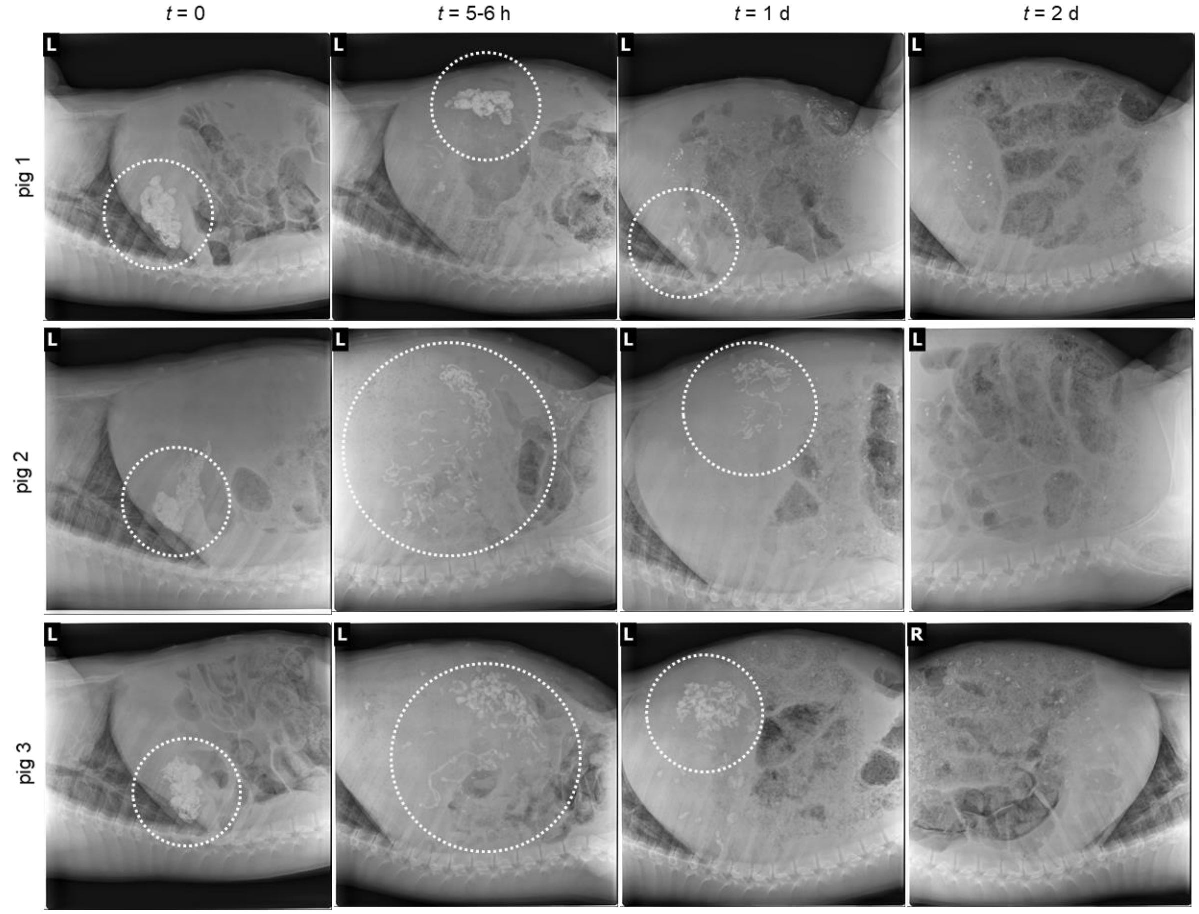


**Figure S6.** *In vivo* retention of LIFT hydrogels. Pigs ($n$ = 3) were administered a solution of crosslinker (200 mM $CaCl_2$/10 mM PEG-dithiol) followed by polymer (0.5% alginate/5% w/v 4-arm PEG-maleimide) loaded with 20% w/v $BaSO_4$ to facilitate X-ray imaging. Hydrogels were present within the stomach up to 24 h after administration.

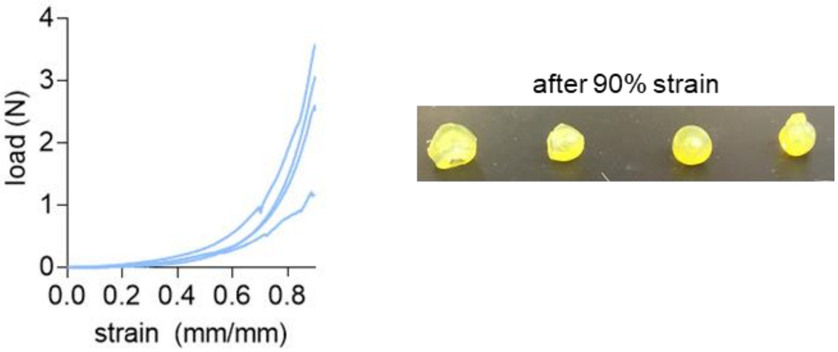


**Figure S7.** Left: load-strain curves of LIFT hydrogels after 90% strain. Hydrogels were formed *in vivo* by administration of a crosslinker solution (200 mM $CaCl_2$/10 mM DMSA) followed by polymer solution (0.5% alginate/5% w/v 4-arm PEG-maleimide). Hydrogels were retrieved 6-8 h after administration. Right: images of LIFT hydrogels after 90% strain. Hydrogels recovered a portion of their initial geometry.

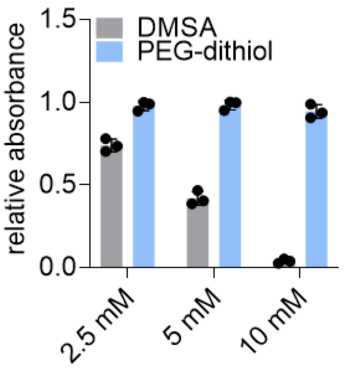


**Figure S8.** Lactase activity after exposure to dithiol compounds. Lactase was added to either DMSA or PEG-dithiol at the indicated concentrations and incubated at 37 °C, 50 RPM, 20 min. Lactase activity was quantified by addition of ONPG and analysis of the colored product. Absorbance was normalized to an untreated lactase control.

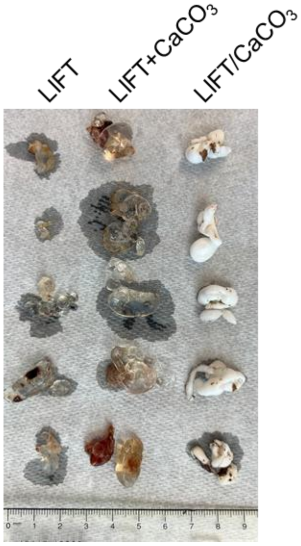


**Figure S9.** LIFT hydrogels after formation in rats. Rats were orally gavaged with a crosslinker solution (200 mM $CaCl_2$/10 mM PEG-dithiol) followed by a polymer solution (0.5% alginate/5% w/v 4-arm PEG-maleimide) with or without $CaCO_3$.

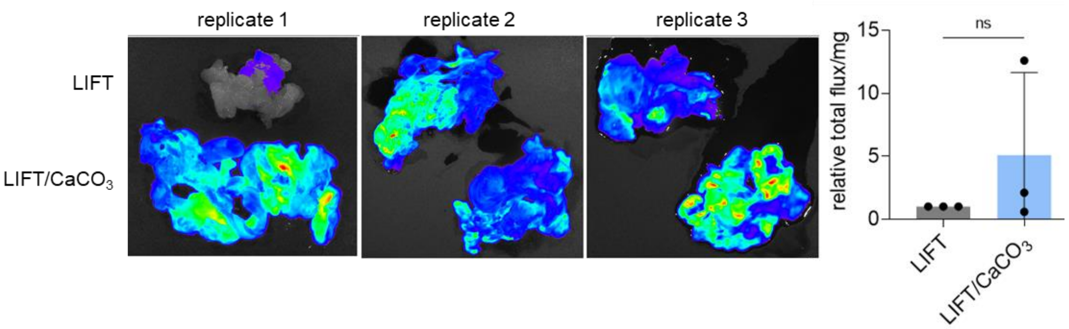


**Figure S10.** Bioluminescence of bacteria encapsulated in LIFT hydrogels with and without $CaCO_3$. Pigs ($n$ = 3) were administered a solution of crosslinker (200 mM $CaCl_2$/10 mM PEG-dithiol) followed by polymer (0.5% alginate/5% w/v 4-arm PEG-maleimide) loaded with $1.6\times10^{10}$ CFU luciferase-expressing bacteria. After 6-8 h, hydrogels were retrieved and bioluminescence was measured, normalized to hydrogel mass, and compared between the two treatments.
